# Erythropoietin production in embryonic neural cells is controlled by hypoxia-inducible factors and histone deacetylases in an undifferentiated state

**DOI:** 10.1101/2024.02.28.582479

**Authors:** Yuma Iwamura, Taku Nakai, Koichiro Kato, Hirotaka Ishioka, Masayuki Yamamoto, Ikuo Hirano, Norio Suzuki

## Abstract

During mammalian development, production sites of the erythroid growth factor erythropoietin (EPO) shift from the neural tissues to the liver in embryos and to the adult kidneys. Embryonic neural EPO-producing (NEP) cells, a subpopulation of neuroepithelial and neural crest cells, express the *Epo* gene between embryonic day (E) 8.5 and E11.5 to promote primitive erythropoiesis in mice. While *Epo* gene expression in the liver and kidney is induced under hypoxic conditions through hypoxia-inducible transcription factor (HIF) 2α, the *Epo* gene regulatory mechanisms in NEP cells remain to be elucidated. This study confirms the presence of cells coexpressing the genes encoding EPO and HIF2α in E9.5 neural tubes, where the hypoxic microenvironment activates HIF1α. In human neural progenitors and mouse embryonic neural tissues, a HIF-activating compound upregulated *EPO* expression, and this induction was blocked by inhibiting HIFs. Additionally, a cell line of NEP cell derivatives that no longer expressed the *Epo* gene demonstrated that histone deacetylase inhibitors (HDACIs) reactivate EPO production while rejuvenating the cells. HDACIs also induced *EPO* gene expression in SK-N-BE(2)c human neuroblastoma cells and mouse primary neural crest cells. Thus, EPO production is controlled by epigenetic mechanisms and hypoxia signaling in an immature state of hypoxic NEP cells.

## Introduction

For maintenance of the oxygen supply to organs in adult mammals, red blood cell production (erythropoiesis) is strictly regulated by the erythroid growth factor erythropoietin (EPO), which is secreted by the kidneys in response to oxygen-depleted conditions (hypoxia) (Suzuki and Yamamoto 2016; Semenza 2023). Since EPO is essential for both primitive and definitive erythropoiesis in fetuses as well as adult erythropoiesis, EPO deficiency results in defects in the differentiation, maturation, and proliferation of erythrocytes in each developmental stage (Wu et al. 1995; Malik et al. 2013; Suzuki et al. 2013; Yamazaki et al. 2013). In developing mouse embryos, the sites of EPO production are shifted from neural tissues to livers around embryonic day (E) 10.5 (Palis 2014; Hirano and Suzuki 2019). Intriguingly, erythropoietic sites are also switched from yolk sac and fetal liver to adult bone marrow during development along with the shift in EPO-producing sites. We and others have demonstrated that hypoxia-inducible factor 2α (HIF2α) activates *EPO* gene transcription in the liver and kidney (Gruber et al. 2007; Tojo et al. 2015; Kobayashi et al. 2016; Souma et al. 2016). However, little is known about the *EPO*-gene regulatory mechanism and contribution of hypoxic signaling to EPO production in neural tissues of midgestation stage embryos, which are hypoxic due to growing organs and immature vascular networks (Lee et al. 2001).

In the kidney, a subset of fibroblasts (renal EPO-producing [REP] cells) localized in the peritubular interstitia secrete EPO to stimulate adult erythropoiesis in the bone marrow (Eckardt et al. 1992; Obara et al. 2008; Pan et al. 2011; Suzuki and Yamamoto 2016; Kragesteen et al. 2023). Because kidney disease conditions inactivate HIF2α in REP cells, chronic kidney disease patients often suffer from renal anemia with EPO deficiency (Sato et al. 2019a). To treat renal anemia, recombinant EPO reagents have been used for more than 30 years. Recently, compounds inhibiting prolyl hydroxylase domain proteins (PHDs), which are negative regulators of HIFs, have been launched as a new class of medicines for renal anemia (Nakai et al. 2023). In addition to REP cells, hepatocytes supportively produce EPO in adult mammals, whereas hepatocytes are the major EPO-production site in late-stage embryos and neonates until the kidneys initiate EPO production (Suzuki et al. 2011; Hirano et al. 2017; Yamazaki et al. 2013,2021).

Before the initiation of hepatic EPO production in embryos, neural EPO-producing (NEP) cells in the neuroepithelium and neural crest secrete EPO (Suzuki et al. 2013; Hirano and Suzuki 2019). As renal and hepatic EPO are required for erythropoiesis in adult bone marrow and fetal liver, respectively, EPO production in NEP cells is also needed for normal primitive erythropoiesis in the yolk sac and bloodstream (Malik et al. 2013; Suzuki et al. 2013). During mouse development, the first *Epo* gene expression is detectable around neural folds by E8.5 at the latest, and EPO is stacked around NEP cells until initiation of circulation (Suzuki et al. 2013). Then, maturation of the circulation system comprising vasculogenesis, angiogenesis and cardiac development makes EPO delivery to erythroid precursors in the yolk sac possible at approximately E9.0, followed by EPO-dependent erythroid maturation. At approximately E9.5, hepatoblasts begin EPO production to induce fetal liver erythropoiesis, whereas NEP cells cease EPO production. Thus, EPO production in neural tissues is transient in midgestation stage embryos during development (Suzuki et al. 2013; Hirano and Suzuki 2019).

While HIF1α and HIF2α are master regulators of cellular and systemic responses to hypoxia, both are required for the development of various organs, including the nervous system. HIF1α-deficient mice exhibit lethality around E10.5 with abnormalities in neural tube formation and neural crest maturation (Iyer et al. 1998; Compernolle et al. 2003). HIF2α deficiency also causes neural abnormalities and embryonic lethality in mice (Tian et al. 1998; Compernolle et al. 2002; Scortegagna et al. 2003). The necessity of HIFs in normal development implies that the hypoxic environment or oxygen gradient in fetuses (Morriss and New 1979) influences morphogenesis and cell differentiation by regulating the stability and activity of HIFs.

Cell-type specific HIF2α-knockout mice demonstrated that HIF2α, but not HIF1α, is required for basal and inducible *Epo* gene expression in hepatocytes after E14.5 and in REP cells (Tojo et al. 2015; Souma et al. 2016; Suzuki et al. 2024). Additionally, we have previously demonstrated that HIF2α induces *Epo* gene expression in REP cells and hepatocytes through the far upstream genomic region from the transcription start site and the proximal downstream region of the transcription end site, respectively (Suzuki et al. 2011; Yamazaki et al. 2013,2021). These findings led us to speculate that *Epo* gene expression in NEP cells, which are a subset of neuroepithelial and neural crest cells in midgestation stage embryos, is also induced by hypoxia in a HIF-dependent manner.

Epigenetic regulatory mechanisms play crucial roles in gene regulation widely related to cell fate determination under physiological and pathological conditions. For instance, histone modification and DNA methylation contribute to the development of the neuroepithelium and neural crest (Martins-Taylor et al. 2012; Hu et al. 2012; Jacob et al. 2014; Rao and LaBonne 2018). Additionally, we and others have reported that nucleosomes are repositioned in the *EPO* gene promoter for hepatic EPO production (Tojo et al. 2015) and that an increase in DNA methylation in the *Epo* gene promoter of REP cells is one of the causes of renal EPO deficiency in renal anemia patients (Yin and Blanchard 2000; Chang et al. 2016; Sato et al. 2019a). Moreover, epigenetic regulators are known to control the fate of neural cells. In particular, histone deacetylases (HDACs) regulate the differentiation of neural lineage cells by suppressing the expression of genes critical for cellular immaturity (Murko et al. 2013; Jacob et al. 2014; Rao and LaBonne 2018). Therefore, epigenetic mechanisms may control *EPO* gene expression in NEP cells during neural development.

The goal of this study was to elucidate the mechanisms of *EPO* gene expression in NEP cells, which is essential for normal primitive erythropoiesis. Because erythrocyte circulation is essential for oxygen delivery to developing organs, the embryo-specific hypoxic microenvironment is considered a trigger that drives EPO production in NEP cells. Then, we investigated the roles of hypoxia signaling and epigenetic mechanisms in *EPO* gene regulation in NEP cells from humans and mice.

## Results

### Hypoxia signaling is active in NEP cells

Since our previous report discovered NEP cells to be murine embryonic neural cells by expressing fluorescent protein transgenes under the control of *Epo* gene regulation (Suzuki et al. 2013), this study began with the detection of endogenous *Epo* gene expression in embryonic neural cells by means of *in situ* hybridization (ISH). In agreement with *Epo* reporter expression, endogenous *Epo* mRNA was exclusively detected in a subset of cells in the neural tube and liver of E9.5 mouse embryos (Fig. 1A). Neural *Epo* expression decreased with development until E13.5, while hepatic expression was sustained, as the fetal liver is known to produce EPO in late-stage embryos (Fig. 1A,B). To identify NEP cells in a species other than mice, ISH of sections from rat embryos at the stage corresponding to murine E9.5 (E11.5) was conducted. As expected, *Epo* mRNA-expressing cells were detected in the rat neural tube (Fig. 1C), indicating that a subset of embryonic neural cells, which has been defined as NEP cells, commonly produces EPO at the midgestation stages in rodents.

**Figure 1.**
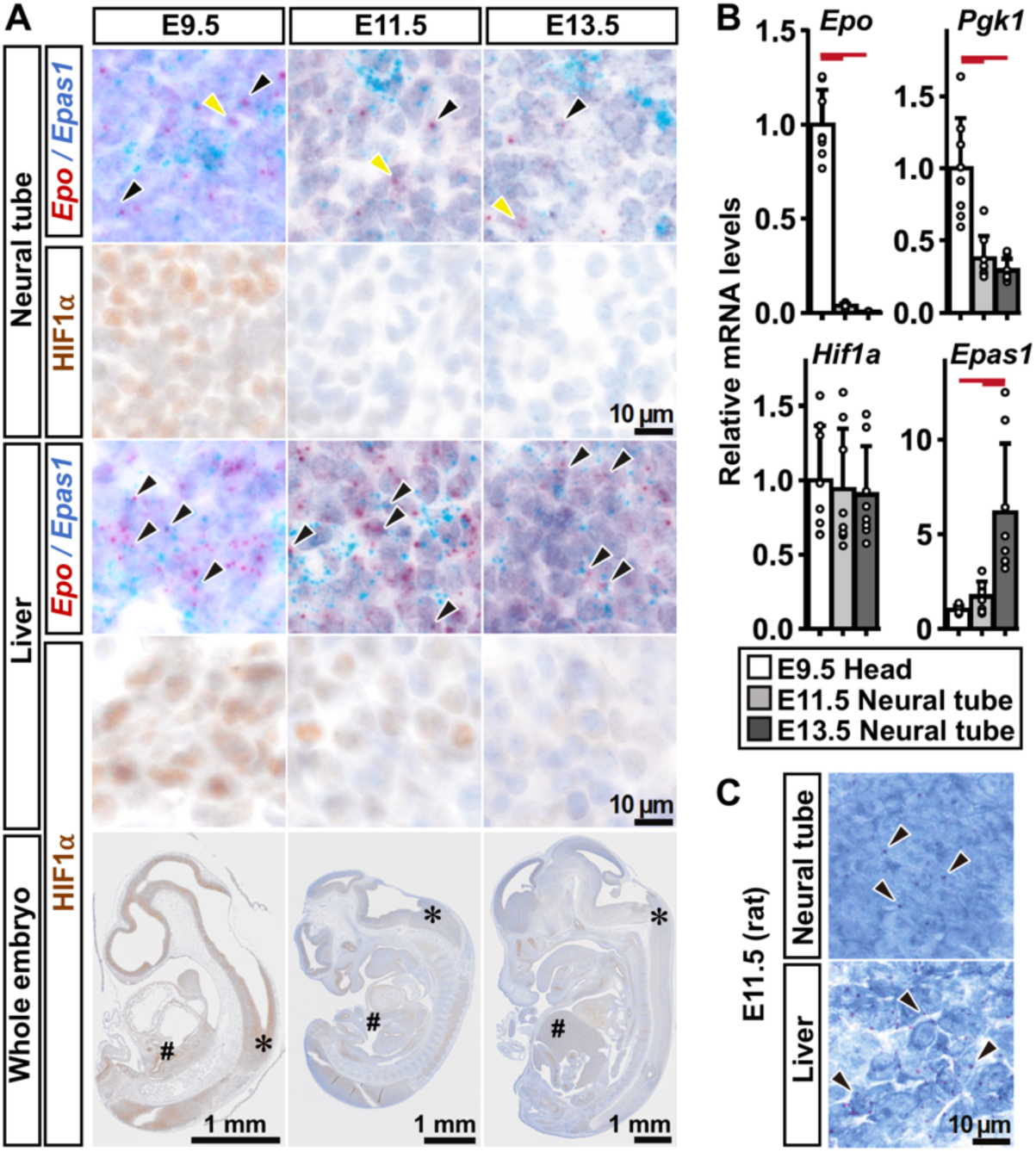
NEP cells in neural tubes express *Epas1* mRNA and HIF1α. (*A*) Representative images of ISH detecting *Epo* (red) and *Epas1* (blue) mRNAs in the neural tube and fetal liver of mouse embryos at E9.5, E11.5 and E13.5. Serial sections of each ISH image were stained with anti-HIF1α antibody (brown). Representative images of HIF1α-immunostained sections from the whole embryo proper are also shown to indicate the neural tubes (*) and livers (#) exhibited in the upper panels. Note that there are NEP cells positive (black arrowheads) and negative (yellow arrowheads) for *Epas1* expression. (*B*) Relative mRNA expression levels of the indicated genes in the E9.5 embryonic head regions, E11.5 neural tubes, and E13.5 neural tubes. The expression levels of E9.5 head samples were set as 1.0 in each graph. n = 6–8 per group. Data are shown as the mean ± standard deviation. Red lines indicate P < 0.01 in the Tukey‒Kramer HSD test. (*C*) Representative images of ISH detecting *Epo* mRNA (brown, arrowhead) in the neural tube and liver of E11.5 rat embryos. Hematoxylin (purple) was used for counterstaining (*A* and *C*).

Next, to examine whether the PHD/HIF pathway is involved in EPO production in NEP cells, the expression of HIFs was analyzed in normal mouse embryos. Intense accumulation of HIF1α in nuclei was widely detected in the neural tube and liver of E9.5 mouse embryos, whereas it disappeared after E11.5 (Fig. 1A). The neural expression levels of *Hif1a* mRNA were unchanged at E9.5, E11.5 and E13.5 (Fig. 1B), suggesting that the hypoxic microenvironment in E9.5 embryos induces HIF1α nuclear accumulation and that the hypoxic milieu is abrogated by establishment of the oxygen delivery system into embryos during mouse development until E11.5. Accordingly, *Pgk1* (phosphoglycerate kinase 1) gene expression, which is known to be induced by hypoxia through HIF1α (Semenza et al. 1991; Suzuki et al. 2018b), was downregulated in neural tissues after E11.5 (Fig. 1B). Because HIF2α detection in embryonic sections was difficult due to a lack of antibody specificity, mRNA expression of the gene for HIF2α (*Epas1*) was estimated. ISH clearly demonstrated *Epas1* mRNA expression in most NEP cells coexpressing *Epo* mRNA as well as hepatocytes (Fig. 1A). The ISH data showing the existence of NEP cells negative for *Epas1* mRNA expression indicate that NEP cells are heterogeneous and that not all NEP cells require HIF2α to produce EPO. In contrast to the *Epo* expression profile, *Epas1* expression was increased during embryonic development (Fig. 1B), suggesting the expansion of *Epas1*-expressing cells such as endothelial cells in developing embryos (Fig. 1A; Tian et al. 1997).

### HIFs enhance EPO gene expression in human and murine NEP cells

Since we confirmed that the genes for EPO and HIF2α are coexpressed in some cells in embryonic neural tissues, we then examined whether *EPO* gene expression is regulated by HIF2α in NEP cells as well as REP cells and hepatocytes (Tojo et al. 2015; Souma et al. 2016). Reverse transcription-quantitative PCR (RT‒qPCR) of primary-cultured human neural progenitor cells (hNPCs) showed that there are cells expressing the *EPO* gene in human embryos (Fig. 2A). The *EPO* mRNA levels were elevated by a PHD inhibitor (GSK360A), which induced *PGK1* gene expression and accumulation of HIF1α and HIF2α in hNPCs (Fig. 2A,B). *EPO* induction was completely abrogated by a HIF2α-specific inhibitor, PT2385 (Wallace et al. 2016), while *PGK1* induction, which is known to be regulated mainly by HIF1α rather than HIF2α, was unaffected (Fig. 2A). These data demonstrate the existence of NEP cells in humans and the contribution of HIF2α to the hypoxic induction of EPO production in NEP cells.

**Figure 2.**
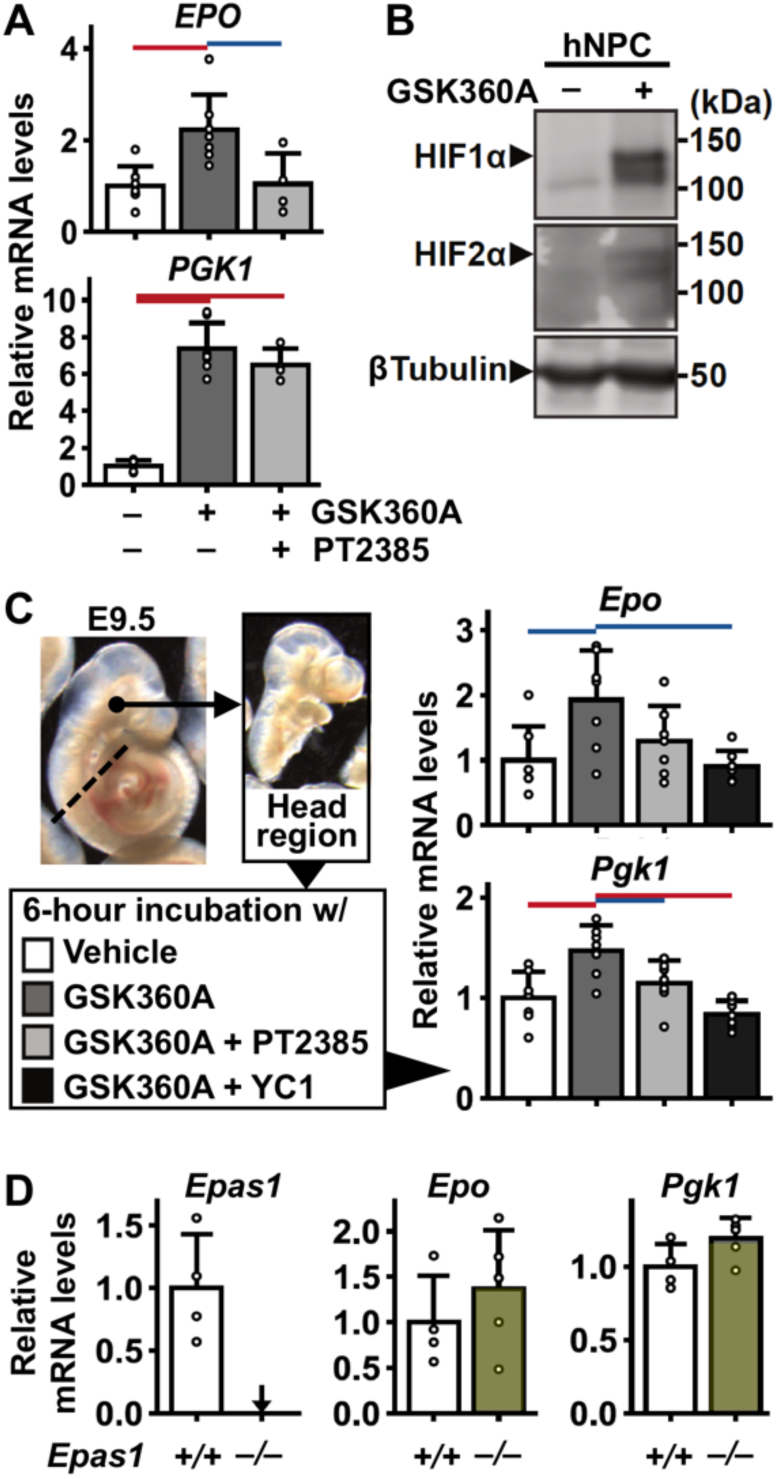
HIFs enhance *EPO* mRNA expression in embryonic neural cells of humans and mice. (*A*) Relative mRNA expression levels of the *EPO* and *PGK1* genes in primary cultured human neural progenitor cells (hNPCs) incubated with PT2385 and/or GSK360A for 24 hours. (*B*) Immunoblots of HIF1α and HIF2α in hNPCs incubated with or without GSK360A for 24 h. βTubulin was used as the loading control. (*C*) Relative mRNA expression levels of the *Epo* and *Pgk1* genes in the head regions from E9.5 mouse embryos that were incubated *ex vivo* with compounds (GSK360A, PT2385 and YC1) affecting HIF activity for 6 hours. (*D*) Relative mRNA expression of the *Epas1, Epo* and *Pgk1* genes in the head regions of *Epas1^+/+^* and *Epas1^−/−^* mouse embryos at E9.5. Expression levels in the vehicle-treated (*A* and *C*) or *Epas1^+/+^* (*D*) samples were set as 1.0 in each graph. Arrows indicate undetectable levels. Data are shown as the mean ± standard deviation. Red and blue lines indicate P < 0.01 and P < 0.05, respectively, in the Tukey‒Kramer HSD test (*A* and *C*) or Student’s *t* test (*D*). n = 6–8 (*A*) and 4–6 (*C* and *D*) per group.

The head regions of E9.5 mouse embryos, in which NEP cells are abundant (Suzuki et al. 2013), were isolated and incubated with compounds influencing the activity of HIFs *ex vivo* for 6 hours to examine the responses of *Epo* gene expression in NEP cells to HIF activity (Fig. 2C). In addition to the hypoxia-inducible *Pgk1* gene, the basal expression levels of the *Epo* gene were enhanced by GSK360A (Fig. 2C), suggesting that HIF signaling upregulates EPO production in NEP cells. HIF2α-specific inhibition by PT2385 slightly decreased the induction, but the reduction was insignificant (Fig. 2C). YC1, an inhibitor of both HIF1α and HIF2α (Li et al. 2008), was then added to the *ex vivo* incubation of head regions with GSK360A. Significant suppression of GSK360A-mediated *Epo* mRNA induction was observed (Fig. 2C). These data suggest that HIF1α and HIF2α redundantly upregulate EPO production in murine NEP cells in response to hypoxia signaling through the hypoxic sensor PHDs.

To investigate the necessity of HIFs in EPO production in NEP cells, *Epo* mRNA levels in the E9.5 head regions of HIF-deficient mice were analyzed. While HIF1α deficiency causes embryonic lethality around E9.5 with abnormal neural development, a previous report demonstrated that HIF1α deficiency results in an approximately 70% reduction in *Epo* mRNA levels in the embryo proper, which contains not only NEP cells but also *Epo*-expressing hepatocytes (Yoon et al. 2006). Our data from E9.5 HIF2α-knockout embryos showed that HIF2α deficiency affected neither *Epo* nor *Pgk1* mRNA levels in NEP cells (Fig. 2D). Thus, it is proposed that HIF1α and HIF2α complementarily activate *Epo* gene expression in NEP cells. In contrast, *EPO* gene induction largely depends on HIF2α expression but not HIF1α expression in hNPCs. This discrepancy in the relevance of HIFs to EPO regulation in NEP cells is likely associated with a small specific fraction of NEP cells in heterogeneous primary cultures of hNPCs. Additionally, basal expression levels of the *Epo* gene are relatively large compared to the HIF-mediated induction levels for EPO production in NEP cells. This profile is different from that of renal and hepatic EPO production, in which marginal basal expression is dramatically increased by hypoxic stimuli.

### HDACIs restore EPO production in NEP cell derivatives

EPO production levels in embryonic neural tissues decreased during development and became very low or undetectable until E11.5 (Fig. 1B; Suzuki et al. 2013). Additionally, NEP cells were identified as immature neural crest/neural progenitors expressing the *Nes* (Nestin), *Foxd3*, *Sox2* and *Wnt1* genes (Suzuki et al. 2013). These previous findings suggest that the differentiation of NEP cells into NEP cell-derived differentiated cells (hereafter “NEP-derived cells”) likely loses EPO production ability along with cellular differentiation/maturation. To test this hypothesis, we analyzed the Neplic cell line as a subpopulation of NEP-derived cells, which we previously established from NEP-derived cells no longer expressing the *Epo* gene in an E15.5 mouse embryo (Hirano and Suzuki 2019). Indeed, *Epo* mRNA expression was undetectable in Neplic cells even in the presence of GSK360A (Fig. 3A).

**Figure 3.**
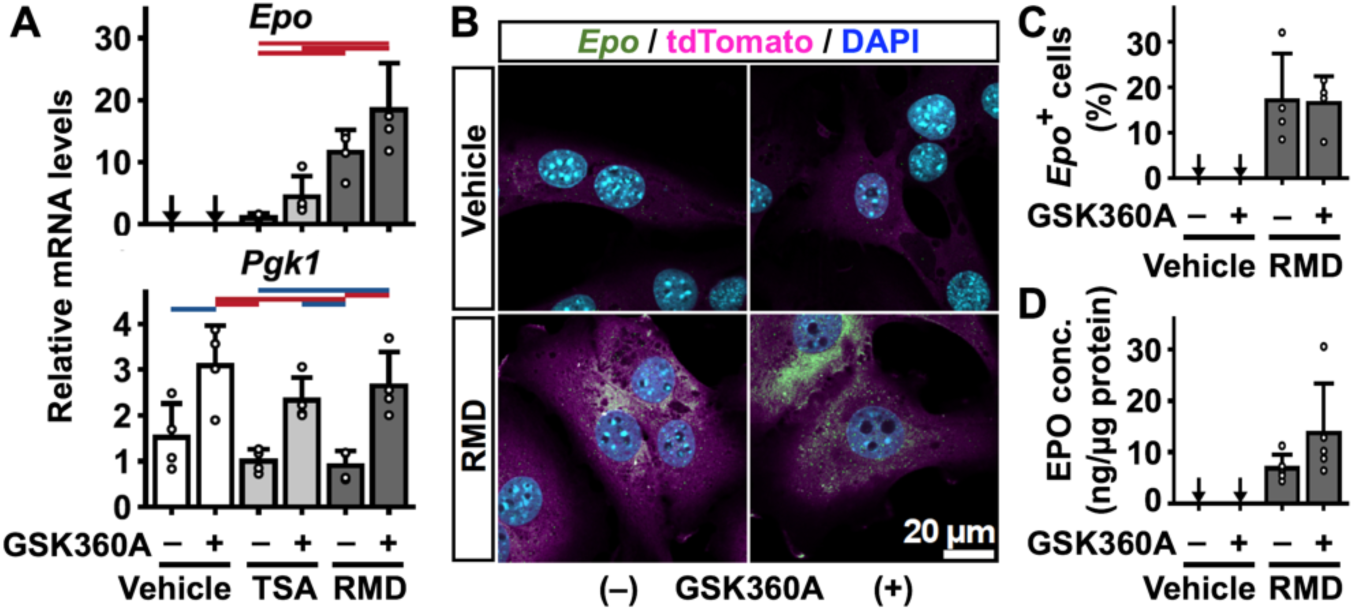
HDACIs restore basal EPO production in NEP-derived cells. (*A*) Relative mRNA expression of the *Epo* and *Pgk1* genes in Neplic cells cultured with TSA, RMD or GSK360A for 24 hours. The expression levels of the TSA-treated samples were set as 1.0. (*B*) Representative images of the ISH experiments detecting *Epo* mRNA (green) in Neplic cells cultured with RMD and/or GSK360A for 24 hours are shown with tdTomato (magenta) and DAPI (cyan) fluorescence. (*C*) Neplic cells positive for *Epo* mRNA expression (*Epo*^+^ cells) were counted in the images from 4 independent ISH experiments (*B*), and the percentages of *Epo*^+^ cells were calculated. (*D*) EPO concentrations in the culture medium of Neplic cells incubated with RMD and/or GSK360A for 24 hours. All data are shown as the mean ± standard deviation. Arrows indicate undetectable levels. n = 3 or 4 (*A*) and 5 (*D*) per group. Red and blue lines indicate P < 0.01 and P < 0.05, respectively, in the Tukey‒Kramer HSD test.

Histone deacetylases (HDACs) are essential for the differentiation and maturation of neural cells, and HDAC inhibitors (HDACIs) potentially induce the rejuvenation of differentiated neural cells to neural progenitors (Montgomery et al. 2009; Jacob et al. 2014). We therefore examined whether HDACIs restore EPO production by inducing the rejuvenation of NEP-derived cells to NEP cells. As expected, both HDACIs, trichostatin A (TSA) and romidepsin (RMD), induced *Epo* mRNA expression in Neplic cells (Fig. 3A). The induction levels were higher in the cells treated with 1 µM RMD than in those treated with 1 µM TSA. PHD inhibition slightly enhanced HDACI-mediated *Epo* mRNA induction in Neplic cells, but the increase was insignificant in contrast to *Pgk1* mRNA induction, which is a known HIF target gene (Fig. 3A). In addition to TSA and RMD, another HDACI, sodium butyrate, also restored *Epo* mRNA expression in Neplic cells (Supplemental Fig. S1; Ajamian et al. 2004). Because RMD most highly induced *Epo* gene expression in Neplic cells among the HDACIs tested, further experiments in this study mainly used RMD.

*Epo* mRNA induction in Neplic cells incubated with RMD was also detected by fluorescence ISH (Fig. 3B,C), which indicated that approximately 20% of Neplic cells became apparently positive for *Epo* mRNA expression after RMD supplementation regardless of the presence of GSK360A. The data from an enzyme-linked immunosorbent assay (ELISA) demonstrated that RMD induces EPO secretion from Neplic cells (Fig. 3D). Additionally, in agreement with the *Epo* mRNA expression profile, HIF activation slightly enhanced the amount of EPO protein secretion (Fig. 3D).

### HDACIs induce histone acetylation and cellular rejuvenation in Neplic cells

Whereas the impacts of HIF activation were marginal for HDACI-mediated *Epo* induction in Neplic cells, nuclear accumulation of HIF1α was dramatically induced by GSK360A (Fig. 4A). The accumulation was slightly lowered by RMD in accordance with previous reports demonstrating attenuation of HIF activity by HDACIs (Mie Lee et al. 2003). Summarizing the data from mouse embryos and Neplic cells, HIFs may supportively contribute to *Epo* gene expression in NEP cells, which is largely controlled by the cellular undifferentiated state.

**Figure 4.**
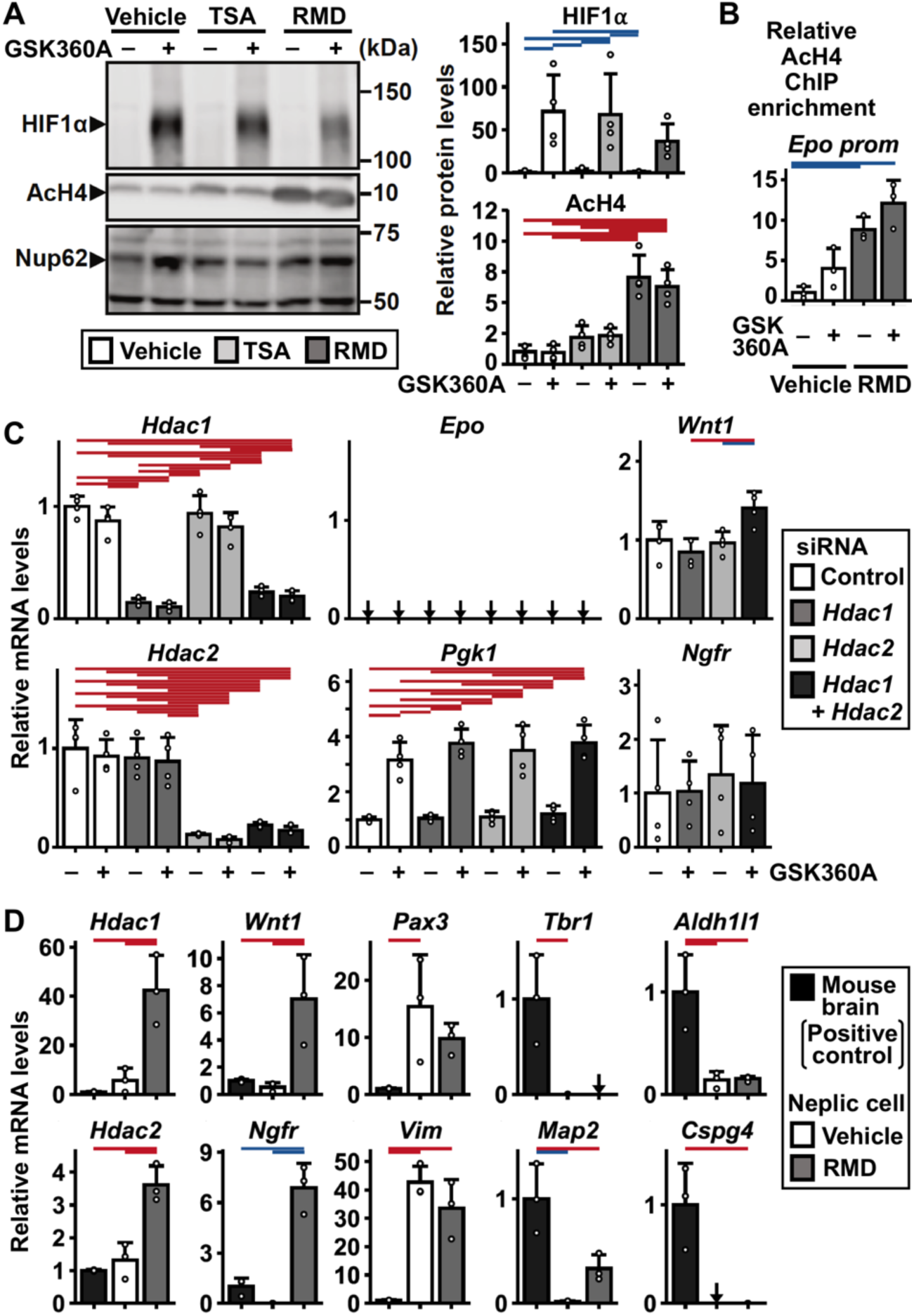
HDAC inhibition induces the rejuvenation of Neplic cells. (*A*) Immunoblots of HIF1α and acetylated histone H4 (AcH4) in the nuclear extracts of Neplic cells cultured with TSA, RMD and/or GSK360A for 24 hours (left). Nup62 was used as the loading control. Relative protein levels of HIF1α and AcH4 were quantified from repetitive immunoblotting after normalization to Nup62 levels (right). The levels in the cells treated without any compounds were set as 1.0 for each graph. (*B*) ChIP‒qPCR analyses with anti-AcH4 antibody on the *Epo* promoter region of Neplic cells incubated with RMD and/or GSK360A for 24 hours. (*C*) Relative mRNA expression of the indicated genes in Neplic cells, in which *Hdac1* and/or *Hdac2* mRNA levels were knocked down by siRNA transfection. The expression levels in the control scramble siRNA transfectants were set as 1.0 for each gene. (*D*) RT‒qPCR analyses of the indicated genes related to neural development in Neplic cells incubated with or without RMD for 24 hours. Expression levels in the adult mouse brain were set as 1.0 for each gene expression. All data are shown as the mean ± standard deviation. Arrows indicate undetectable levels. n = 3 or 4 per group. Red and blue lines indicate P < 0.01 and P < 0.05, respectively, in the Tukey‒ Kramer HSD test.

TSA and RMD were confirmed to increase nuclear concentrations of acetylated histone H4 (AcH4) in Neplic cells (Fig. 4A). Chromatin immunoprecipitation analyses (ChIP) with anti-AcH4 antibody demonstrated that RMD significantly increases acetylated histones around the *Epo* gene promoter in Neplic cells (Fig. 4B). Consistent with the *Epo* mRNA expression profile, AcH4 levels in the *Epo* promoter were slightly elevated after HIF activation. These results suggest that HDACIs reactivate *Epo* gene expression in Neplic cells *via* their canonical effect, which increases histone acetylation by inhibiting HDACs.

Since we demonstrated that three HDACIs commonly activate *Epo* gene expression in Neplic cells, probably through enhancing histone acetylation, the relevance of each HDAC isoform to *Epo* gene induction was investigated by conducting knockdown experiments using small interfering RNA (siRNA). Knocking down the expression of HDAC1 and/or HDAC2, both of which are major targets of RMD, did not recapitulate the effect of HDACIs on *Epo* gene induction in Neplic cells (Fig. 4C). These data suggested that the knockdown levels are insufficient in this study or that HDAC isoforms other than HDAC1 and HDAC2 are involved in HDACI-mediated *Epo* induction.

The effects of HDACIs on the differentiation state of Neplic cells were examined by RT‒qPCR of neural cell marker genes after incubation of the cells with RMD for 24 hours. Interestingly, RMD significantly induced the expression of the *Hdac1* and *Hdac2* genes (Fig. 4D), suggesting a compensatory induction mechanism to maintain cellular HDAC activity (Zupkovitz et al. 2006). The expression of the *Wnt1* and *Ngfr* genes, which are expressed in NEP cells and neural progenitors (Suzuki et al. 2013), was induced in Neplic cells after supplementation with RMD or sodium butyrate (Fig. 4D; Supplemental Fig. S1). These data suggested that HDACIs induce the rejuvenation of NEP-derived cells into neural progenitor-like NEP cells. Accordingly, the *Vim* and *Pax3* genes, which are also known to be expressed in neural progenitors (Schnitzer et al. 1981; Goulding et al. 1991), were highly expressed in Neplic cells cultured with or without RMD (Fig. 4D). On the other hand, the expression levels of neuronal (*Tbr1* and *Map2*) and glial (*Aldh1l1* and *Cspg4*) marker genes remained low after RMD treatment compared to those in the adult mouse brain (Fig. 4D; De Camilli et al. 1984; Neymeyer, et al. 1997; Ong and Levine 1999; Hevner et al. 2001). Consistent with the results regarding *Epo* gene induction, siRNA-mediated knockdown of *Hdac1* and *Hdac2* expression did not affect the expression of neural cell lineage markers, except for a slight but statistically significant induction of *Wnt1* gene expression in *Hdac1/2* double knockdown cells (Fig. 4C). These results indicate that Neplic cells are considered neural lineage cells that differentiate from NEP cells partially maintaining their neural progenitor signatures, and HDAC inhibition restores the EPO production ability, which is lost during NEP cell differentiation, by inducing the rejuvenation of Neplic cells into NEP cell-like neural progenitors.

### HDAC inhibition induces EPO gene expression in human neuroblastoma cells

We verified the effects of HDACIs on neural cells other than Neplic cells. RMD supplementation adequately recapitulated *EPO* gene induction in the SK-N-BE(2)c cell line, which is derived from human neuroblastoma (Fig. 5A; Suzuki et al. 2018b). As expected, AcH4 levels in the *EPO* promoter were elevated by RMD (Fig. 5B). The effects of HIF activation by GSK360A on the mRNA and AcH4 levels of the *EPO* gene were minor (Fig. 5A,B). Additionally, the expression levels of neural progenitor marker genes (*WNT1* and *NGFR*) were dramatically induced by RMD in SK-N-BE(2)c cells (Fig. 5C). Changes in gene expression profiles related to mature neural cells (*TBR1*, *MAP2*, *ALDH1L1* and *CSPG4*) were insignificant between with and without 24-hour RMD treatment (Fig. 5C). These data from a human cell line suggest that HDAC inhibition induces EPO production and rejuvenation not only in Neplic cells but also in other types of mammalian neural lineage cells.

**Figure 5.**
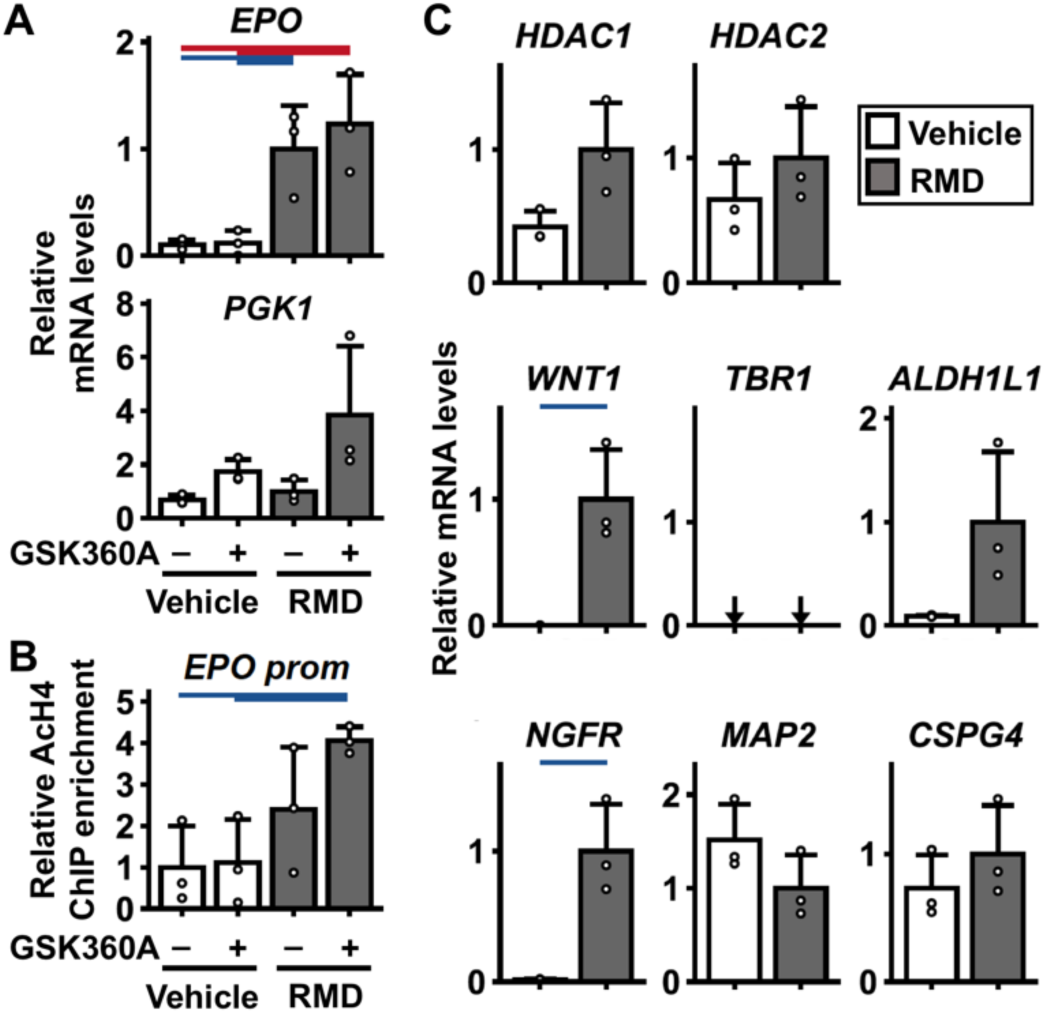
HDACI-mediated *EPO* gene induction is reproduced in human neuroblastoma cells. (*A*) Relative mRNA expression of the *EPO* and *PGK1* genes in SK-N-BE(2)c cells incubated with RMD and/or GSK360A for 24 hours. The expression levels of the vehicle-treated samples were set as 1.0 for each gene. (*B*) ChIP‒qPCR analyses with anti-AcH4 antibody on the *EPO* promoter region of SK-N-BE(2)c cells incubated with RMD and/or GSK360A for 24 hours. ChIP enrichment in the vehicle-treated samples was set as 1.0. (*C*) Relative mRNA expression of genes related to histone deacetylation (*HDAC1* and *HDAC2*), neural progenitor markers (*NGFR* and *WNT1*), neuron markers (*TBR1* and *MAP2*) and glial markers (*ALDH1L1* and *CSPG4*) in SK-N-BE(2)c cells incubated with or without RMD for 24 hours. Expression levels in the RMD-treated samples were set as 1.0 for each gene expression. Data are shown as the mean ± standard deviation. n = 3 per group. Arrows indicate undetectable levels. Red and blue lines indicate P < 0.01 and P < 0.05, respectively, in the Tukey‒Kramer HSD test (*A* and B) or Student’s *t* test (*C*).

### HDAC inhibition reactivates Epo gene expression in primary neural crest cells

We further validated that HDACIs induce EPO production and rejuvenation in primarily cultured murine neural crest cells containing NEP-derived cells. Since our previous report demonstrated that *Epo* gene expression is mainly detected in the cardiac neural crest (Suzuki et al. 2013), the neural tubes of cardiac regions were isolated from E9.5 mouse embryos and incubated *ex vivo* (Fig. 6A; Etchevers 2011). We confirmed that the isolated neural tube and the migrating neural crest cells from them in the Day-1 primary culture system contained NEP cells or NEP-derived cells, which were labeled with tdTomato fluorescence in the REC mouse line (Fig. 6B; Yamazaki et al. 2013; Suzuki and Yamamoto 2016).

**Figure 6.**
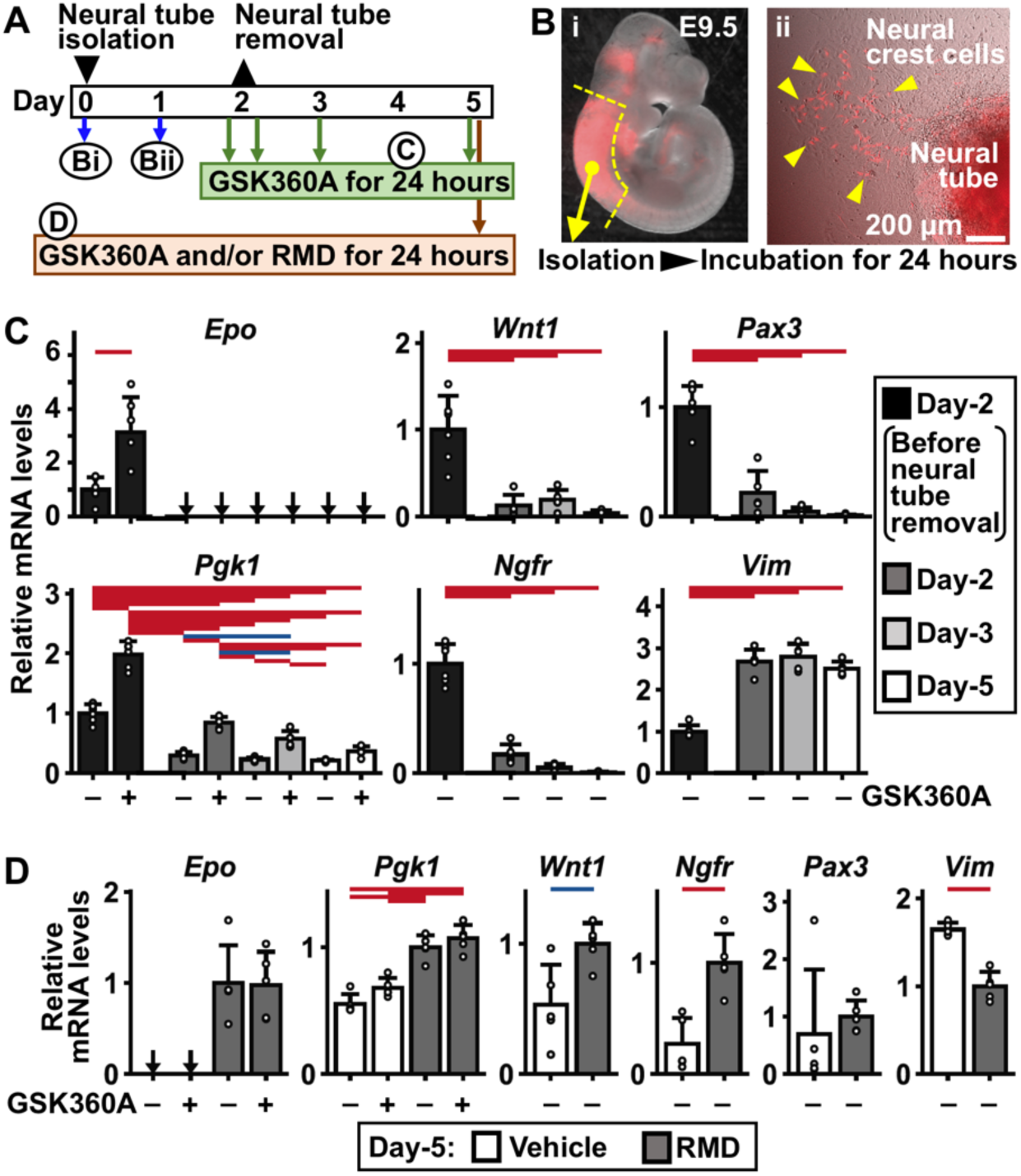
HDACIs recall *Epo* expression in primary neural crest cells. (*A*) An experimental scheme for the primary culture of neural crest cells from the neural tubes of E9.5 mouse embryos. On Day-2, lumps of the neural tubes were removed from the culture, and the remaining neural crest cells were cultured for 3 days. B, C and D indicate the samples for the data shown in panels *B*, *C* and *D*, respectively. (*B*) Representative tdTomato fluorescence images, which indicate NEP cells or NEP-derived cells, were taken from the E9.5 REC embryo (i) or the primary culture system on Day-1 (ii). Yellow arrowheads indicate cells positive for tdTomato migrating out from the neural tube (ii). (*C*) Relative mRNA expression of the indicated genes in the primary neural crest cells incubated with GSK360A for 24 hours. The primary neural crest cells were taken on Day-2 before and after neural tube removal as well as on Day-3 and Day-5 (*A*). (*D*) Relative mRNA expression of the indicated genes in Day-5 primary neural crest cells incubated with RMD and/or GSK360A for 24 hours. All data are shown as the mean ± standard deviation (*C* and *D*). n = 5 per group. Arrows indicate undetectable levels. Red and blue lines indicate P < 0.01 and P < 0.05, respectively, in the Tukey‒Kramer HSD test.

*Epo* mRNA expression was detected in the primary culture system containing whole neural tube cells primarily incubated *ex vivo* for 2 days and was significantly enhanced by GSK360A-mediated HIF activation (Fig. 6C). In contrast, in neural crest cells isolated from the primary culture system by removing the neural tubes, *Epo* mRNA expression was not detected even in the presence of GSK360A (Fig. 6C). The basal and induction levels of *Pgk1* gene expression in response to GSK360A gradually decreased during culturing *ex vivo* (Fig. 6C), suggesting that HIF activity is attenuated in the culture system. Since the expression levels of 3 of 4 neural progenitor marker genes examined were lowered in the neural crest cells during incubation compared to those of the Day-2 primary culture system just before removal of the neural tubes (Fig. 6C), cells migrating out of the neural tubes contained differentiated neural crest cells. These data indicated that embryonic neural cells bearing EPO production ability, which are NEP cells, localize and remain around the neural tube of mouse embryos at E9.5. Additionally, NEP cells were considered to lose the EPO-production ability while differentiating into migrating neural crest cells out of the neural tube.

Since the primary culture system of neural crest cells containing NEP-derived cells was established, we then investigated the impacts of HDAC inhibition on *Epo* gene expression and cellular rejuvenation. Consistent with the data from Neplic and SK-N-BE(2)c cells, RMD supplementation induced *Epo* gene expression in primary neural crest cells (Fig. 6D). Additionally, the expression of the neural progenitor markers *Wnt1* and *Ngfr* was also elevated, while that of the other neural progenitor markers, *Pax3* and *Vim*, was not altered, as in Neplic and SK-N-BE(2)c cells. These results supported our contention that HDAC inhibition restores EPO-production ability, which is lost during differentiation of NEP cells, through inducing rejuvenation of NEP-derived cells.

### Hypoxia allows rejuvenated NEP-derived cells to retain the EPO production ability

Since this study proposed HIF-inducible *Epo* gene expression in NEP cells located in the hypoxic microenvironment of developing mouse embryos, we then tested the relevance of the hypoxic environment to the EPO production ability and/or cellular undifferentiated state of NEP cells. The basal and HIF-inducible *Epo* gene expression in Neplic cells treated with RMD disappeared 3 days after washing out RMD from the culture medium, while *Pgk1* gene expression maintained HIF responsiveness regardless of the presence of RMD (Fig. 7A,B). *Epo* gene induction in response to HIF activation was examined in Neplic cells that were incubated under hypoxic (1% oxygen) conditions after RMD treatment. The RT–qPCR data showed that hypoxia sustains HIF-inducible, but not basal, *Epo* gene expression, which is restored by HDAC inhibition, in NEP-derived cells (Fig. 7A,B).

**Figure 7.**
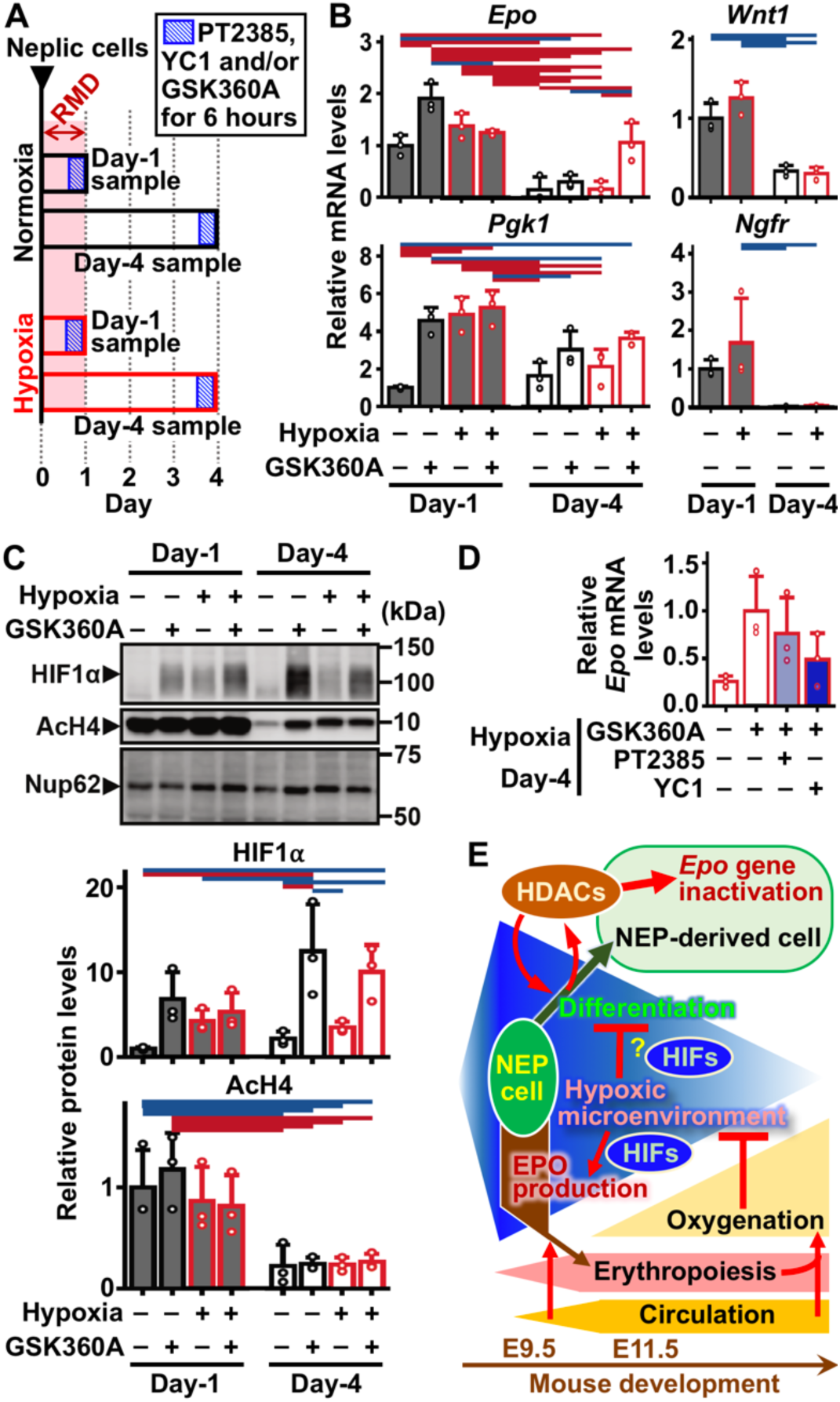
HDACI-treated NEP-derived cells retain the EPO production ability under hypoxic conditions. (*A*) A scheme of an experiment to determine the impacts of hypoxia on RMD-treated Neplic cells. RMD was added to Neplic cells for 24 hours under normoxic or hypoxic (1% oxygen) conditions and washed out on Day-1. The cells were incubated with GSK360A and/or PT2385 or YC1 for 6 hours before (Day-1 samples) or after (Day-4 samples) 72-hour continuous incubation under normoxic or hypoxic conditions. (*B*) Relative mRNA expression of the indicated genes in Day-1 and Day-4 samples cultured in normoxia or hypoxia with or without GSK360A. (*C*) Immunoblots of HIF1α and AcH4 in the nuclear extracts of in Day-1 and Day-4 samples cultured in normoxia or hypoxia with or without GSK360A. Nup62 was used as the loading control (upper). Relative protein levels of HIF1α and AcH4 were quantified from repetitive immunoblotting after normalization to Nup62 levels (lower). (*D*) Relative mRNA expression of the *Epo* gene in Day-4 samples cultured in hypoxia with or without GSK360A, PT2385 and YC1 for 6 hours. All data are shown as the mean ± standard deviation. n = 3 per group. Red and blue lines indicate P < 0.01 and P < 0.05, respectively, in the Tukey‒Kramer HSD test. The levels in the Day-1 samples cultured in normoxia without GSK360A were set as 1.0 for each graph (*B, C* and *D*). (*E*) Schematic representation of the regulatory mechanism of EPO production along with NEP cell differentiation in developing mouse embryos. The intense blue area indicates hypoxic microenvironment around NEP cells, which affects differentiation and EPO production in NEP cells.

Induced expression of neural progenitor markers (*Wnt1* and *Ngfr*) and elevated levels of AcH4 in RMD-treated Neplic cells were decreased after removing RMD under both normoxic and hypoxic conditions (Fig. 7B,C). These data indicate that HDAC inhibition induces rejuvenation of NEP-derived cells transiently and reversibly. Additionally, hypoxia is insufficient to maintain the RMD-induced immature state of Neplic cells. Hypoxia-sustained *Epo* gene inductivity was marginally attenuated by either PT2385 or YC1 (Fig. 7D). Consequently, the hypoxic microenvironment around NEP cells seems to be involved in the maintenance of EPO production in resistance to HDAC-mediated *Epo* gene suppression during development (Fig. 7E).

## Discussion

EPO production in NEP cells located in the neural crest and neuroepithelium is essential for normal primitive erythropoiesis (Suzuki et al. 2013; Hirano and Suzuki 2019). The current study elucidated the regulatory mechanism of EPO production in NEP cells, which is associated with hypoxia signaling and the cellular undifferentiated state with epigenetic modifications (Fig. 7E). These results provide critical information on the molecular mechanisms of erythropoietic regulation during mammalian development.

NEP cells are developmentally the first site of EPO production, which was discovered in transgenic mice expressing GFP reporter transgenes under the control of the mouse *Epo* gene regulatory system (Suzuki et al. 2013). In the current study, we adopted an innovative ISH technique and successfully detected endogenous *Epo* gene expression in the neural tissues of mouse and rat embryos. Additionally, analyses of human fetus-derived neural progenitors (hNPCs) demonstrated the existence of NEP cells in humans. Moreover, EPO production has been reported in the larval brains of frogs and fish (Paffett-Lugassy et al. 2007; Nogawa-Kosaka et al. 2010). These findings suggest that NEP cells, which are neural cells that produce EPO to promote primitive erythropoiesis, are conserved among vertebrates. EPO deficiency in mouse embryos and fish larvae leads to no apparent abnormalities in organs unrelated to erythropoiesis (Suzuki et al. 2002, 2013; Paffett-Lugassy et al. 2007). However, the current study does not exclude the possibility that the function of EPO in NEP cells goes beyond erythropoiesis, since many reports have demonstrated the non-erythropoietic functions of EPO, such as neuroprotection (Yu et al. 2002; Jia et al. 2012; Constanthin et al. 2020).

NEP cells are the major site of EPO production in midgestation stage embryos (Suzuki et al. 2013; Hirano and Suzuki 2019). In the other major EPO-producing cells (hepatocytes in late embryos and REP cells in adults), EPO production levels largely depend on hypoxia signaling to maintain the oxygen supply to organs through red blood cells (Suzuki and Yamamoto 2016; Suzuki et al. 2024). In contrast, this study demonstrates that EPO production levels in NEP cells are highly dependent on the cellular differentiation state through epigenetic mechanisms and that HIF activation adds slight induction levels to basal level expression. Additionally, the low-level hypoxic induction of *Epo* gene expression in NEP cells is controlled redundantly by HIF1α and HIF2α, while HIF2α, but not HIF1α, dramatically induces *Epo* gene expression in hepatocytes and REP cells (Tojo et al. 2015; Souma et al. 2016). Thus, the regulatory mechanism of EPO production in NEP cells is different from that in hepatocytes and REP cells.

The enhancer regions for *Epo* gene expression in REP cells and hepatocytes are located upstream and downstream of the gene coding region, respectively (Suzuki et al. 2011; Hirano et al. 2017). Our previous studies using reporter transgenic mice demonstrated that neither of the enhancer regions are needed for *Epo* gene expression in NEP cells (Suzuki et al. 2013; Hirano et al. 2017). Thus, in addition to HIF usage and hypoxic induction, *cis*-regulatory elements of the *Epo* gene are different among the 3 major EPO production sites. Since this study developed a strategy to reactivate *Epo* gene expression in NEP-derived cells, chromatin and epigenetic analyses using *Epo*-reactivated cells are anticipated to elucidate the molecular mechanisms of *Epo* gene regulation in NEP cells.

Even though the HIF-dependent induction of EPO production in NEP cells is weak compared to that in the other EPO production sites, the induction indicates that primitive erythropoiesis is controlled in a hypoxia-inducible manner to maintain oxygen homeostasis as well as definitive erythropoiesis (Suzuki and Yamamoto 2016). In agreement with previous reports showing an embryo-specific hypoxic milieu before establishment of the stable circulation system (Lee et al. 2001; Makita et al. 2001), our data demonstrate that HIF1α accumulation is induced by tissue hypoxia in the neural tube around NEP cells and disappears by oxygenation after initiation of circulation. Consequently, *Epo* gene expression in the 3 distinct cell types responsible for EPO production in each life stage is commonly regulated by the oxygen microenvironment around the cells to maintain systemic oxygen homeostasis (Fig. 7E). Furthermore, it is possible that oxygen distribution in developing embryos is somewhat involved in boosting the initiation of primitive erythropoiesis by enhancing EPO production in NEP cells.

NEP cells are found in the neural crest and neuroepithelium of midgestation stage embryos and disappear after E11.5 in mice (Suzuki et al. 2013). Tracing the fate of NEP cells in genetically modified mice, NEP-derived cells have been widely identified in various organs, including the central nervous system. (Yamazaki et al. 2013; Hirano and Suzuki 2019). Therefore, NEP cells are immature cells that are mainly differentiated into neural lineage cells that lose EPO production ability during development. This study elucidated that the cellular undifferentiated state is associated with the EPO-production ability of NEP cells using NEP-derived ‘Neplic cells’ incubated with HDACIs, which have been reported to induce rejuvenation of some neural cells *via* changes in epigenetic modifications (Shakèd et al. 2008; Montgomery et al. 2009). Because 3 types of HDACIs commonly induce *Epo* gene expression and cellular rejuvenation in Neplic cells, HDACs are thought to play central roles in the differentiation and EPO production of NEP cells (Fig. 7E). Indeed, histone acetylation levels in the *EPO* promoter increase in Neplic and SK-N-BE(2)c cells after HDACI treatment. However, further studies using Neplic cells are needed to elucidate the molecular mechanisms by which HDAC isoform(s) govern EPO production and differentiation in NEP cells.

EPO production in NEP cells is inactivated, probably due to epigenetic mechanisms related to cellular differentiation into NEP-derived cells. Likewise, the EPO production ability of REP cells in adult kidneys is impaired due to epigenetic mechanisms, including DNA methylation, which are associated with transformation from fibroblasts to myofibroblasts under pathological conditions (Sato et al. 2019ab). Moreover, the *Epo* gene regulatory mechanism in hepatocytes is altered during cellular differentiation from hepatoblasts to hepatocytes in fetuses, and gene expression is attenuated to negligible levels in adult hepatocytes even under hypoxic conditions (Suzuki et al. 2011; Yamazaki et al. 2021; Nakai et al. 2023). These findings suggest that epigenetic mechanisms associated with the cellular immature/undifferentiated state are closely related to the acquisition and retention of EPO production ability (Fig. 7E).

EPO production levels in NEP cells are more likely related to the cellular undifferentiated state than oxygen conditions, in contrast to those in hepatocytes and REP cells. Although the regulatory mechanism of *Epo* gene expression in NEP cells is different from that in REP cells, the idea that a subset of NEP cells is differentiated into REP cells while retaining or reactivating EPO production ability is of interest (Hirano and Suzuki 2019; Tsujimoto et al. 2024). In fact, REP cells in the adult kidney, which exhibit a part of the gene expression profiles of neural cells, are reported to originate from the neural crest (Obara et al. 2008; Asada et al. 2011; Kragesteen et al. 2023). Regarding the fate of NEP cells, particularly their relevance to REP cells, further studies using mouse models and additional cell lines are anticipated to elucidate the development of the mammalian erythropoietic system. In addition, we propose that adult neural cells are potential therapeutic targets of EPO induction for EPO-deficiency anemia in chronic kidney disease patients.

## Materials and Methods

### Animals

All animal experiments were approved by the Animal Care Committee of Tohoku University (2020MdA-078-03 and 2018MdA-164-03). Pregnant Slc:SD rats and ICR mice were purchased from Japan SLC to isolate embryos (Shizuoka, Japan). *Epas1^+/-^* mice with a C57black/6 background were generated by deleting *Epas1* exon 2 in the germline of *Epas1* conditional knockout mice, which were purchased from the Jackson Laboratory (Gruber et al. 2007). To obtain *Epas1^-/-^* embryos, *Epas1^+/-^* parents were crossed, and 12:00 PM on the day a vaginal plug was observed was denoted E0.5. The REC mouse line (*Rosa26^+/LSL-tdTomato^:Tg^EpoCre^*genotype), in which cells that have experienced *Epo* gene expression are permanently labeled with tdTomato fluorescence beyond mitosis, was described previously (Yamazaki et al. 2013; Suzuki and Yamamoto 2016). To determine the genotypes of the embryos, genomic DNA from yolk sacs was used for PCR with previously described primers (Yamazaki et al. 2013; Tojo et al. 2015).

### Histological analyses

Tissue sections (5 µm thickness) of paraffin-embedded mouse embryos, which were fixed in 4% paraformaldehyde (Nacalai Tesque, Kyoto) at room temperature overnight, were retrieved using a retriever (R2100, Aptum Biologics). For blocks of small tissues, E9.5 embryos were embedded in iPGel (Genostaff, Tokyo) before fixation. Chemogenic ISH was performed according to the instructions of RNAscope 2.5 HD Duplex Assay kit or Brown Assay kit with probes specific for *Epo* and *Epas1* mRNAs (Advanced Cell Diagnostics) (Miyauchi et al. 2021). For immunohistochemistry, sections were incubated with diluted HIF1α antibody (Abnova) at 4 °C overnight after incubation with Blocking One Histo (Nacalai Tesque), and then signals (brown color) were developed with the ImmPRESS Excel Amplified Polymer Staining kit (Vector Laboratories). Mayer’s hematoxylin solution (Merck) was used for counterstaining. For fluorescent ISH of cultured cells, cells fixed with 4% paraformaldehyde at room temperature for 1 hour were stained with a probe set specific for mouse *Epo* mRNA using the ISH Palette system (Nepagene, Tokyo; Tsuneoka and Funato 2020). tdTomato, which is constitutively expressed in Neplic cells, and 4′,6-diamidino-2-phenylindole (DAPI, Sigma) were used for counterstaining. Cells containing more than 4 dots of *Epo* mRNA signals were counted as *Epo*^+^ cells after the image processing. Images were obtained using a VS200 research slide scanner (Olympus), an m165 stereomicroscope (Leica), a BZ-X800 fluorescence microscope (Keyence), or a LSM780 confocal microscope (Zeiss). The details and dilutions of antibodies used in this study are listed in Supplemental Table S1.

### RT‒qPCR

Total RNA was purified using the RNeasy kit (Qiagen) or ISOGEN reagent (Nippongene, Tokyo) for cultured cells. cDNAs were synthesized from the total RNA using Superscript IV reverse transcriptase with random primers (Thermo Fisher Scientific). Quantitative PCR (qPCR) was performed using a LightCycler 96 with SYBR Green (Roche) and gene-specific primers (Supplemental Table S2). Relative mRNA levels were calculated by the 2^−ΔΔCt^ method after normalization to the expression levels of housekeeping genes (mouse *Hprt* or human *HPRT* genes).

### Cell culture

Human neural progenitor cells (hNPCs, PT-2599, Lonza), which were derived from a fetus aborted at 16 weeks of pregnancy, were maintained in neuron culture medium (Wako, Osaka) containing 10% fetal bovine serum (FBS, Biosera, Cholet, France). Mouse NEP cell-lineage immortalized and cultivable (Neplic) cells (Hirano and Suzuki 2019) and SK-N-BE(2)c human neuroblastoma cells (Suzuki et al. 2018b) were cultured in Dulbecco’s modified Eagle medium (DMEM) containing 4.5 g/L glucose, antibiotics (Nacalai Tesque), and 10% FBS. For the primary culture of neural crest cells (Etchevers 2011), the cardiac neural tube regions of E9.5 mouse embryos were isolated, and each tissue was incubated with neural crest culture medium containing 10% KnockOut serum replacement (Thermo Fisher Scientific) in each well of 6-well plates coated with collagen I (Corning). After a 48-hour incubation of the neural tubes, the lumps of the neural tube were removed from the culture system, and the remaining neural crest cells that migrated out of the neural tubes were cultured for 3 days. Hypoxic cell culture was performed in a hypoxic workstation (Sci-tive Dual, Baker Ruskinn, Stanford, ME), in which oxygen concentrations were set at 1.0% by nitrogen gas replacement.

### Inhibitors

GSK360A (Toronto Research Chemicals), YC1 (Sigma‒Aldrich), PT2385 (Cayman Chemicals, Ann Arbor), and RMD (FK228, Cayman Chemicals), all of which were dissolved in dimethyl sulfoxide (DMSO, Nacalai Tesque), were used to inhibit PHDs, HIFs, HIF2α and class I HDACs, respectively (Tojo et al. 2015; Suzuki et al. 2018ab; Sato et al. 2019a). For inhibition of class I and II HDACs, trichostatin A (TSA, Cayman Chemicals) and sodium butyrate (Nacalai Tesque) were also used. The final concentrations were 50 µM for GSK360A, 100 µM for YC1, 50 µM for PT2385, and 1 µM for RMD.

### ex vivo incubation of mouse embryonic tissues

Embryo propers at E9.5 were separated into the head and body regions within 1 hour (Fig. 2C), followed by incubation with inhibitors in DMEM supplemented with 10% FBS for 6 hours in a CO_2_ (5%) incubator at 37 °C.

### Immunoblot

Whole cell lysates were obtained from cells lysed with RIPA buffer (50 mM Tris-HCl pH 8.0, 150 mM NaCl, 0.5% sodium deoxycholate, 0.1% SDS, and 1.0% NP-40) containing 10 mM MG132 and 1.0% protease inhibitor cocktail (Nacalai Tesque). For preparation of nuclear extracts, the precipitates of cells lysed with Buffer A (10 mM HEPES pH 7.9, 1.5 mM MgCl_2_, 10 mM KCl, 0.5 mM DTT, 0.05% NP-40, 10 mM MG132, 1.0% protease inhibitor cocktail) were isolated by centrifugation at 3,000 rpm for 10 minutes and resuspended in RIPA buffer (Suzuki et al. 2018b). Protein samples were loaded for electrophoresis in 5% or 15% polyacrylamide gels containing sodium dodecyl sulfate (SDS‒PAGE) and transferred onto nitrocellulose membranes using the Trans-Blot Turbo system (Bio-Rad). The membranes were incubated with the primary antibodies listed in Supplemental Table S1 at 4 °C overnight after blocking with Blocking One reagent (Nacalai Tesque), followed by incubation with HRP-conjugated secondary antibodies (Supplemental Table S1) at room temperature for 1 hour. Chemiluminescent signals were detected with ECL Prime Western blotting detection reagent (Cytiva, Marlborough, MA) using a C-DiGit scanner and quantified using ImageStudio software (LI-COR, Lincoln, NE). βTubulin and Nup62 protein levels were used as loading controls for whole cell extract and nuclear extract, respectively. The details and dilutions of antibodies used in this study are listed in Supplemental Table S1.

### ChIP‒qPCR

ChIP was performed using a ChIP-IT Express Enzymatic kit (Active Motif) following the manufacturer’s instructions. Briefly, chromatin samples cross-linked by formalin were enzymatically shared and incubated with rabbit anti-AcH4 antibody or isotype control antibody (Merck) at 4 °C overnight. Antibody-binding chromatin fragments were purified with protein G magnetic beads. After reverse crosslinking, samples were subjected to qPCR with primers (Supplemental Table S2). To calculate relative ChIP enrichment, the inputs of each sample were used.

### EPO ELISA

EPO concentrations in the culture medium of Neplic cells incubated with RMD for 24 hours, which were concentrated with Amicon Ultra Centrifugal Filters (Merck), were measured for the concentration using a Mouse EPO ELISA Kit (R&D) and for the total protein level using the Protein Assay CBB solution (Nacalai Tesque).

### siRNA knockdown

The siRNA smart pool against murine *Hdac1* and *Hdac2* (25 pmol/well, Horizon Discovery, Cambridge) and scramble siRNA were transfected into Neplic cells cultured in 12-well plates using Lipofectamine 3000 reagent (Invitrogen). The medium was not replaced, and the cells were harvested 48 hours after transfection.

### Statistics

Statistical significance was determined by the Tukey‒Kramer test or Student’s *t* test for comparison of multiple groups or two groups, respectively. The data were recognized as statistically significant at P < 0.05.

## Competing Interest Statement

The authors declare no competing interests.

## Acknowledgments

This study was supported in part by MEXT/JSPS KAKENHI (21H02676, 22K193960 and 23KK0137), the Kidney Foundation Japan (JKFB23-1), the Astellas Foundation for Research on Metabolic Disorders, the Naito Foundation, the Uehara Memorial Foundation, and the Takeda Science Foundation (to NS) as well as the JST SPRING (JPMJSP2114 to YI). The funders had no role in this study. We thank the Pathology section of the Biomedical Research Core of Tohoku University Graduate School of Medicine, Atsuko Konuma and Yuito Sasaki (Tohoku University) for technical help as well as the Biomedical Research Core and Centre for Laboratory Animal Research of Tohoku University.

## Author Contributions

YI and NS developed the project and designed the study. YI, TN, KK, HI and NS performed the experiments and analyzed the data. YI and NS constructed the figures and wrote the manuscript. YI, TN, KK and NS provided animal models and cells. MY, IH and NS supervised the project. All authors discussed the results and suggested revisions.

